# Integrated post-genomic cell wall analysis reveals floating biofilm formation associated with high expression of flocculins in the pathogen *Candida krusei*

**DOI:** 10.1101/2023.01.26.525814

**Authors:** María Alvarado, Jesús Alberto Gómez-Navajas, María Teresa Blázquez-Muñoz, Emilia Gómez-Molero, Carmen Berbegal, Elena Eraso, Gertjan Kramer, Piet W.J. De Groot

## Abstract

The pathogenic yeast *Candida krusei* is more distantly related to *Candida albicans* than clinically relevant CTG-clade *Candida* species. Its cell wall, a dynamic organelle that is the first point of interaction between pathogen and host, is relatively understudied, and its wall proteome remains unidentified to date. Here, we present an integrated study of the cell wall in *C. krusei*. Our comparative genomic studies and experimental data indicate that the general structure of the cell wall in *C. krusei* is similar to *Saccharomyces cerevisiae* and *C. albicans* and is comprised of β-1,3-glucan, β-1,6-glucan, chitin, and mannoproteins. However, some pronounced differences with *C. albicans* walls were observed, for instance, higher mannan and protein levels and altered protein mannosylation patterns. Further, despite absence of proteins with high sequence similarity to *Candida* adhesins, protein structure modeling identified eleven proteins related to flocculins/adhesins in *S. cerevisiae* or *C. albicans*. To obtain a proteomic comparison of biofilm and planktonic cells, *C. krusei* cells were grown to exponential phase and in static 24-h cultures. Interestingly, the 24-h static cultures of *C. krusei* yielded formation of floating biofilm (flor) rather than adherence to polystyrene at the bottom. The proteomic analysis of both conditions identified a total of 32 cell wall proteins. In line with a possible role in flor formation, increased abundance of flocculins, in particular Flo110, was observed in the floating biofilm compared to exponential cells. This study is the first to provide a detailed description of the cell wall in *C. krusei* including its cell wall proteome, and paves the way for further investigations on the importance of flor formation and flocculins in the pathogenesis of *C. krusei*.

**AUTHOR SUMMARY:** The yeast *Candida krusei* is among the five most prevalent causal agents of candidiasis but its mechanisms underlying pathogenicity have been scarcely studied. This is also true for its cell wall structure, an essential organelle that governs primary host-pathogen interactions and host immune responses. Solid knowledge about cell wall synthesis and dynamics is crucial for the development of novel antifungal strategies against this pathogenic yeast. Here, through a combination of comparative genomics, protein structure modeling, and biochemical and proteomic analysis of purified walls, we present a detailed study of the cell wall composition in *C. krusei* and identify important architectural differences compared to *C. albicans* cell walls. Cell walls of *C. krusei* contain higher mannan and protein levels with altered mannan branching patterns, governed by expansions and reductions in gene families encoding mannosyltransferases. We also show that, in contrast to other *Candida* species, static cultures produce floating biofilms. Comparative wall proteomic studies of these biofilms show increased abundance of flocculins and hydrolytic enzymes, protein classes implicated in biofilm formation and primary host-pathogen interactions leading to tissue colonization. In conclusion, our study uncovers important keys towards a better molecular understanding of the virulence mechanisms of the important pathogen *C. krusei*.

## INTRODUCTION

Candidiasis is one of the most frequent fungal infections in humans. In immunocompromised patients *Candida* infections frequently lead to invasive mycoses or bloodstream infections (candidemia), which are associated with high mortality rates (1). *Candida krusei* is among the most relevant etiological agents of candidiasis and is most often found in patients with hematological malignancies or receiving prolonged azole prophylaxis (2, 3). Infections caused by this organism are of special clinical relevance because of its intrinsic resistance to fluconazole (4).

*C. krusei* is a diploid ascomycete yeast belonging to the family *Pichiaceae* in the order Saccharomycetales. It is only distantly related to *C. albicans* (4, 5); moreover, it does not belong to the *Candida*-CTG clade (6). Genetic studies have characterized this pathogenic yeast as the asexual (anamorph) form of *Pichia kudriavzevii* (teleomorph), however, the different nomenclature for the pathogenic form (*C. krusei*) and the environmental teleomorph (*P. kudriavzevii*) is still maintained (5).

Among the virulence factors described for *C. krusei*, we find similarities with other *Candida* spp., for instance, formation of pseudohyphae that would confer the ability to invade host tissues (7), secretion of phospholipases and proteinases that enhance the yeast ability to colonize host tissues and evade the host immune system (8), phenotypic switching contributing to its adaptation to environmental conditions (9), or formation of monospecies or polymicrobial biofilms (10).

Playing a key role in primary host-pathogen interactions, the fungal cell wall is crucial for all the aforementioned virulence mechanisms (11, 12). In related yeasts such as *Saccharomyces cerevisiae* and *C. albicans*, the inner part of the cell wall has been described as a carbohydrate network mostly composed of the polysaccharides β-1,3-glucan, β-1,6-glucan and chitin, whereas the outer cell wall layer is densely packed with highly glycosylated (mostly by mannosyl residues) glycosylphosphatidylinositol (GPI)-modified proteins that are covalently bound to β-1,6-glucan molecules. These proteins belong to different families and have a manifold of functions including enzymatic activities needed for cell wall synthesis and modification, enzymes using (host) substrates in the surrounding environment (e.g aspartic proteases), proteins involved in surface adhesion and biofilm formation, and others (13). Interestingly, genomic analyses showed enrichment of some GPI protein families (e.g. Hyr/Iff adhesins) in pathogenic *Candida* spp. compared to related non-pathogenic species, suggesting a direct relationship to pathogenicity (14).

GPI-modified adhesins confer yeasts the ability to adhere onto different substrates such as host cells or abiotic surfaces (medical devices, catheters), and thus play an important role in the establishment of infections (15, 16). In addition, the ability to form biofilms by cell-to-cell adhesion renders an infiltrate stable matrix that enhances resistance to antifungal agents (17). GPI proteins in *C. albicans* include three adhesin families: Als (18), Hyr/Iff (19) and Hwp (20). Genomic analyses have shown that these adhesin families are conserved in other *Candida* species of the CTG clade (14). For the non-CTG clade pathogenic yeast *Candida glabrata* similar genomic studies demonstrated that it contains an extraordinarily large number of more than 70 sequences encoding GPI-modified adhesin-like proteins (21-25). Recent structural studies showed that most of these adhesins can be divided into two groups: (i) Epa1-related proteins with shown or presumed host-binding lectin activities and containing N-terminal PA14-like domains similar to flocculins in *S. cerevisiae* (26, 27), and (ii) proteins with β-helix/β- sandwich N-terminal domains for which the substrate ligands are still unidentified although involvement in adherence to polystyrene surfaces was demonstrated for Awp2 (28).

Following adhesion to host cells, *Candida* species secrete enzymes that help to disrupt host membranes and proteins and actively penetrate into tissues (29), as well as improve the efficiency of extracellular nutrient acquisition (30). Three different classes of such secreted/cell wall hydrolases, namely aspartic proteases, phospholipases, and lipases have been described (31). Among the aspartic proteases and phospholipases, some members may be retained into the cell wall through GPI anchoring (14, 32, 33). Expansion of genes encoding aspartic proteases in pathogenic species compared to less pathogenic relatives supports a role for these proteins in the infection process (14, 34).

Thorough knowledge of the cell wall structure and its proteomic composition is crucial to better understand biofilm formation and host-pathogen interactions underlying *Candida* pathogenesis. However, in *C. krusei* this remains poorly studied to date. The aim of this study therefore was to enhance our knowledge of the *C. krusei* cell wall through an integrated approach including a comprehensive genomic inventory of its cell wall biosynthetic machinery, analysis of its cell wall composition and proteome, and surface adhesion and biofilm formation studies. Intriguingly, proteomic analysis of flor formed during static culturing revealed increased expression of a Flo11-like wall protein potentially important for biofilm formation during infection.

## Results

### The cell wall synthetic genetic machinery of *C. krusei*

Here, we present a multidisciplinary study to enhance our knowledge about synthesis and composition of the cell wall in the human pathogenic yeast *C. krusei*. First, an extensive bioinformatic analysis of its cell wall biosynthetic genetic machinery was carried out on the translated genome of reference strain CBS573 using known fungal cell wall biosynthetic genes as search queries. Cell walls of CTG-clade *Candida* species and *S. cerevisiae* are mainly composed of a network of the polysaccharides, β-1,3-glucan, β-1,6-glucan, and chitin, to which a variety of glycoproteins are covalently attached either to β-1,6-glucan through GPI remnants, or to β-1,3-glucan through mild-alkali sensitive linkages (ASL). Synthesis of β-1,3-glucan and chitin is carried out by enzyme complexes located in the plasma membrane. The corresponding genes, as well as those of proteins involved in synthesis of precursor molecules or with regulatory functions, for instance the cell wall integrity pathway, appear mostly conserved in *C. krusei* with similar representation in the genome as described for *C. albicans* and *S. cerevisiae* (Table 1; details in S1 Table) (14). However, the genomic analysis of *C. krusei* revealed reductions in the numbers of genes in glycosyl hydrolase families GH5 (Exg), GH81 (Eng), and GH18 (Cht), implicated in modification or hydrolysis of β-1,3-glucan and chitin, for instance during cytokinesis.

**Table 1.**
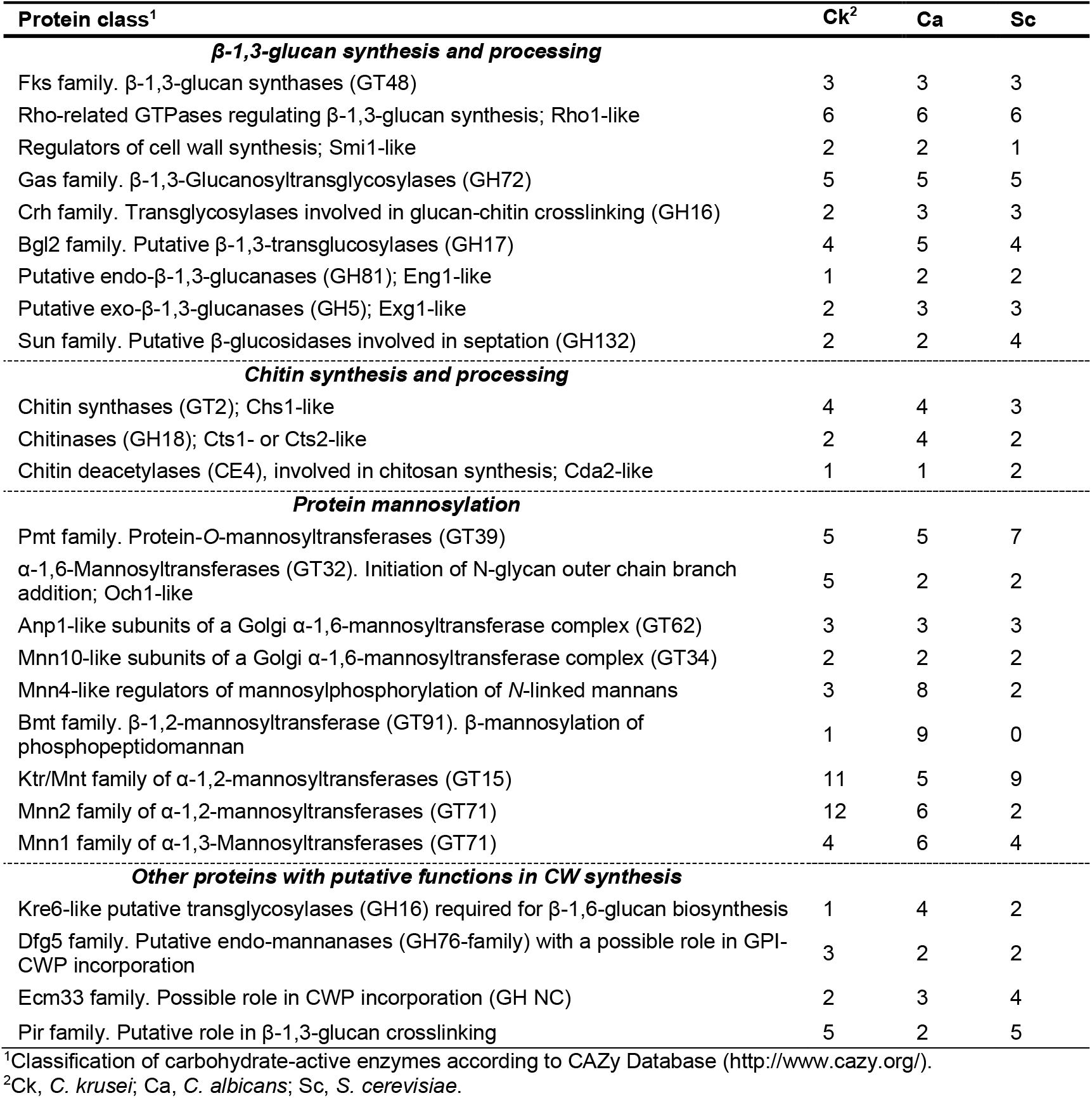
Comparative analysis of protein families involved in cell wall biosynthesis of *C. krusei*.

| Protein class<sup>1</sup> | Ck<sup>2</sup> | Ca | Sc |
| --- | --- | --- | --- |
| ***β-1,3-glucan synthesis and processing*** |  |  |  |
| Fks family. β-1,3-glucan synthases (GT48) | 3 | 3 | 3 |
| Rho-related GTPases regulating β-1,3-glucan synthesis; Rho1-like | 6 | 6 | 6 |
| Regulators of cell wall synthesis; Smi1-like | 2 | 2 | 1 |
| Gas family. β-1,3-Glucanosyltransglycosylases (GH72) | 5 | 5 | 5 |
| Crh family. Transglycosylases involved in glucan-chitin crosslinking (GH16) | 2 | 3 | 3 |
| Bgl2 family. Putative β-1,3-transglucosylases (GH17) | 4 | 5 | 4 |
| Putative endo-β-1,3-glucanases (GH81); Eng1-like | 1 | 2 | 2 |
| Putative exo-β-1,3-glucanases (GH5); Exg1-like | 2 | 3 | 3 |
| Sun family. Putative β-glucosidases involved in septation (GH132) | 2 | 2 | 4 |
| ***Chitin synthesis and processing*** |  |  |  |
| Chitin synthases (GT2); Chs1-like | 4 | 4 | 3 |
| Chitinases (GH18); Cts1- or Cts2-like | 2 | 4 | 2 |
| Chitin deacetylases (CE4), involved in chitosan synthesis; Cda2-like | 1 | 1 | 2 |
| ***Protein mannosylation*** |  |  |  |
| Pmt family. Protein-*O*-mannosyltransferases (GT39) | 5 | 5 | 7 |
| α-1,6-Mannosyltransferases (GT32). Initiation of N-glycan outer chain branch addition; Och1-like | 5 | 2 | 2 |
| Anp1-like subunits of a Golgi α-1,6-mannosyltransferase complex (GT62) | 3 | 3 | 3 |
| Mnn10-like subunits of a Golgi α-1,6-mannosyltransferase complex (GT34) | 2 | 2 | 2 |
| Mnn4-like regulators of mannosylphosphorylation of *N*-linked mannans | 3 | 8 | 2 |
| Bmt family. β-1,2-mannosyltransferase (GT91). β-mannosylation of phosphopeptidomannan | 1 | 9 | 0 |
| Ktr/Mnt family of α-1,2-mannosyltransferases (GT15) | 11 | 5 | 9 |
| Mnn2 family of α-1,2-mannosyltransferases (GT71) | 12 | 6 | 2 |
| Mnn1 family of α-1,3-Mannosyltransferases (GT71) | 4 | 6 | 4 |
| ***Other proteins with putative functions in CW synthesis*** |  |  |  |
| Kre6-like putative transglycosylases (GH16) required for β-1,6-glucan biosynthesis | 1 | 4 | 2 |
| Dfg5 family. Putative endo-mannanases (GH76-family) with a possible role in GPI-CWP incorporation | 3 | 2 | 2 |
| Ecm33 family. Possible role in CWP incorporation (GH NC) | 2 | 3 | 4 |
| Pir family. Putative role in β-1,3-glucan crosslinking | 5 | 2 | 5 |

<sup>1</sup>Classification of carbohydrate-active enzymes according to CAZy Database (http://www.cazy.org/).<sup>2</sup>Ck, *C. krusei*; Ca, *C. albicans*; Sc, *S. cerevisiae*.

Synthesis of cell wall β-1,6-glucan in yeasts is a poorly understood process although the molecule has a crucial role in interconnecting β-1,3-glucan, chitin, and GPI-modified mannoproteins. For instance, a β-1,6-glucan-synthetizing enzyme remains unidentified to date even though Kre6-like GH16 proteins have been postulated as candidates (35, 36). Noteworthy, where *C. albicans* contains four and *S. cerevisiae* two Kre6-like paralogs, Kre6 and its twin paralog Skn1 that arose from the whole genome duplication, *C. krusei* contains only a single Kre6 homolog in its genome.

The Dfg5 GH76 family of putative endo-mannanases is described to hydrolyze glycan moieties of GPI anchors for subsequent attachment of GPI proteins to non-reducing ends of β-1,6-glucan in the cell wall (37-39). Most *Candida* species and *S. cerevisiae* contain two *DFG5*-related genes, but three paralogs are present in *C. krusei* (Table 1). In contrast, for the Ecm33 family, proposed to have a - still unresolved - role in CWP or β-1,6-glucan incorporation (40, 41), only two genes are present in *C. krusei* whereas *C. albicans* contains three and *S. cerevisiae* four (two twin pairs) paralogs.

Proteins that are destined to be covalently bound to the cell wall polysaccharide network are secretory proteins. After synthesis of their precursors, they are translocated to the cell surface and usually become highly glycosylated during their passage through the endoplasmic reticulum and Golgi apparatus. Glycosylation in *Candida* occurs via two different processes, *O*- and *N*-glycosylation. *O*-glycans are short mannan chains of up to five residues connected to hydroxyl side chains of serine (Ser) and threonine (Thr) residues. As CWPs usually have a high percentage of these residues, *O*-glycosylation contributes significantly to the mannan present in the cell wall of *Candida. N-*glycosylation occurs less frequently, as its acceptor molecules are side chain nitrogen atoms of asparagine (Asn) residues that are part of an Asn–X–Ser/Thr consensus sequence (where X is any amino acid except proline). However, individual *N*-glycan chains may contain up to 200 mannose residues, added by diverse families of mannosyltransferases, and thus also greatly contribute to the cell wall mannan content. Interestingly, in *C. krusei* clear differences are observed in the gene copy repertoire of mannosyltransferases compared to *C. albicans*. While *C. albicans* contains two Och1-like GT32 α-1,6-mannosyltransferases needed for *N*-glycan outer chain backbone formation, *C. krusei* contains five Och1 paralogs. Further, the GT15 (Ck 11 vs Ca 5 genes) and GT71 (Ck 12 vs Ca 6 genes) families of presumed α-1,2-mannosyltransferases are also largely extended in *C. krusei*. On the other hand, the analyzed genome of *C. krusei* contains only one GT91 β-1,2-mannosyltransferase gene, whereas this family consists of nine paralogs in *C. albicans*, and the number of Mnn4-like regulators of mannosylphosphorylation of *N*-linked mannans is also reduced (Ck 3 vs Ca 9 genes). The Pmt family of protein-*O*-mannosyltransferases (GT39) comprises five paralogs in both species.

We also identified putative GPI proteins and CWPs covalently bound through mild-alkali sensitive linkages (ASL). Our bioinformatic pipeline selected 43 putative GPI protein candidates (S2 Table), which is less than described for pathogenic CTG-clade *Candida* spp. (14, 19, 42), *C. glabrata* (about 100) (43) and *S. cerevisiae* (about 70) (42) using similar approaches. Nonetheless, the list of predicted *C. krusei* GPI proteins includes most of the protein families also described in the abovementioned species, for example, Gas, Crh, Ecm33, Dfg5, adhesins/flocculins, aspartyl proteases, and phospholipases. Detailed manual inspection of protein sequences identified during mass spectrometric CWP analysis (see below) indicated that some GPI proteins were not picked up (false negatives) by our GPI protein pipeline, probably mostly due to erroneous automated annotations.

Blast searches identified ten probable aspartyl proteases in *C. krusei*, six of which are predicted to be GPI anchored. For comparison, 13 aspartyl proteases have been described in *C. albicans*, two of which are documented as GPI proteins with cell wall localization (33). In *C. glabrata*, nine predicted GPI-modified aspartyl proteases were identified (24) but, consistent with their absence in cell wall preparations, these proteins present dibasic motifs in the region immediately upstream of the GPI attachment site, favoring plasma membrane retention (44).

Eleven ORFs showing adhesin properties and/or weak similarity to Flo1, 5, 9 and 10 and Flo11 flocculins in *S. cerevisiae* or Ywp1/Hwp1 adhesins in *C. albicans* were identified (Table 2 and S2 Table). Despite the low level or even absence of sequence similarity, their identifications as putative flocculins and adhesins was supported by tertiary (3D) structure modeling and subsequent structure-similarity analysis (Fig 1). Remarkably, (part of) the tertiary structures of the three ORFs with weak primary structure similarity to Ywp1 showed more reminiscence, albeit with relatively high RMSD values, to β-sandwich structures encountered in Als3. We tentatively designated these proteins <u>a</u>dhesin-like <u>p</u>roteins 1-3 (Alp1-3; Table 2). Manual inspection of the flocculin ORF sequences revealed that the majority seems to have incorrect ORF boundaries (S1 Table), providing a likely explanation why half of the flocculins were not identified by the GPI prediction pipeline but probably are genuine GPI proteins, in accordance with their presence in cell wall preparations (Tables 2 and S3 Table).

**Table 2.** Identified adhesin/flocculin-like proteins in *C. krusei* strain CBS573.

| NCBI Refseq | Proposed Name | Closest Sc or Ca Blast homolog<sup>1</sup> | Closest PDB homolog (RMSD (Å)) | Identified in cell wall |
| --- | --- | --- | --- | --- |
| XP_029321299 | Flo9 | Flo9 | ScFlo5, 2XJP (2.2) | Yes |
| XP_029321005 + XP_029321006 | Flo90 | Flo9 | ScFlo5, 2XJP (2.6) | Yes |
| XP_029320844 | Flo91 | Flo9 | ScFlo5, 2XJP (2.6) |  |
| XP_029323952 | Flo92 | Flo9 | ScFlo5, 2XJP (2.4) | Yes |
| XP_029322636 | Flo10 | Flo9 | ScFlo5, 2XJP (2.2) | Yes |
| XP_029322875 | Flo11 | Flo11 | ScFlo11, 4UYR (2.3) |  |
| XP_029321007 | Flo110 | no hits | ScFlo11, 4UYR (2.7) | Yes |
| XP_029322143 | Flo111 | no hits | ScFlo11, 4UYR (3.2) | Yes |
| XP_029319891 | Alp1 | Ywp1 | CaAls3, 4LEE (4.5) |  |
| XP_029319890 | Alp2 | Ywp1 | CaAls3, 4LEE (4.6) |  |
| XP_029323142 | Alp3 | Ywp1 | CaAls3, 4LEE (6.8) |  |

<sup>1</sup>Sc, *S. cerevisiae*; Ca, *C. albicans*.

**Figure 1.**
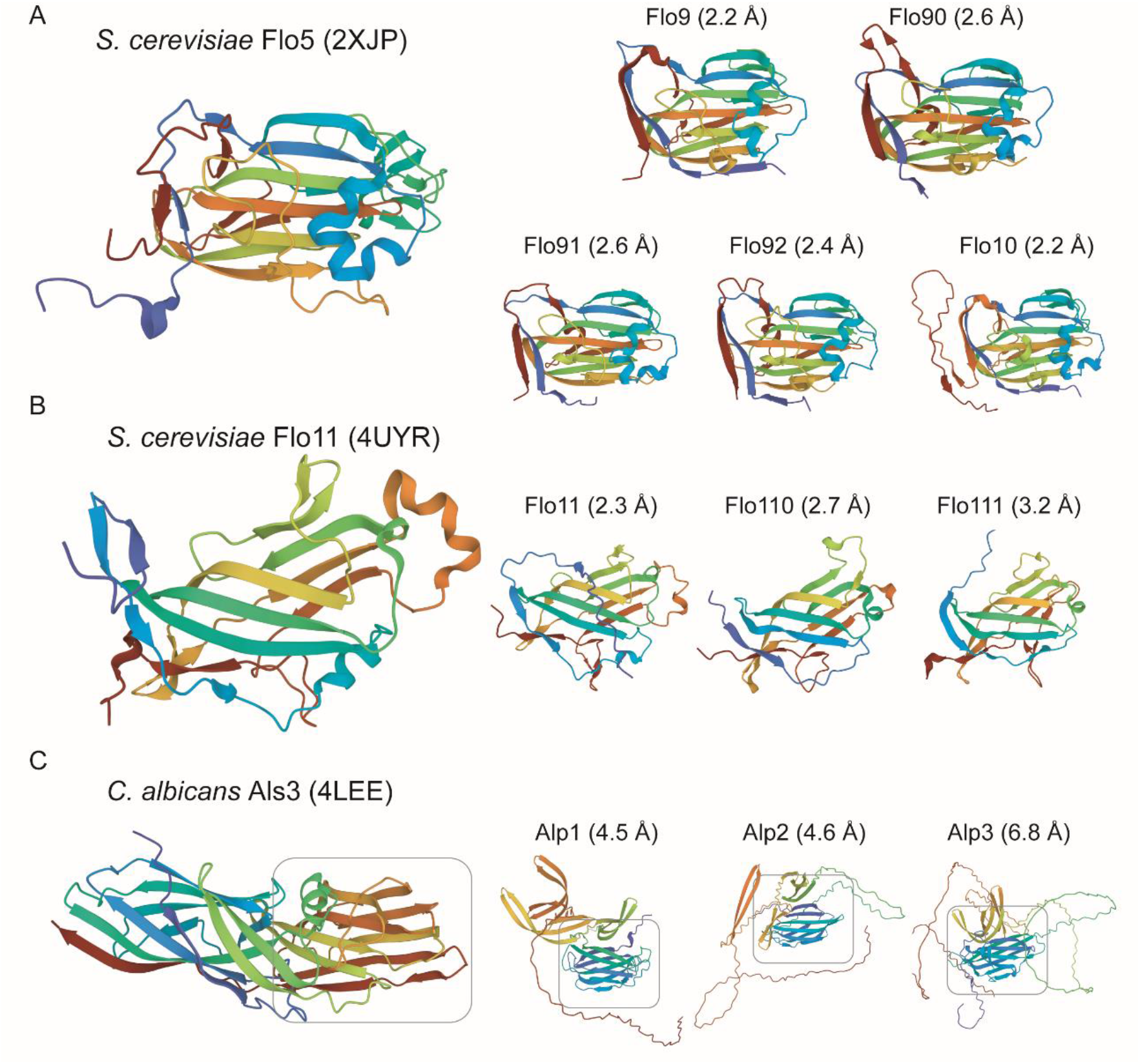
Tertiary (3D) structure analysis of *C. krusei* adhesins by AlphaFold modeling of putative ligand-binding domains. Structures are shown in rainbow-color representation from blue (N-terminus) to red (C-terminus). *C. krusei* ORFs with similarity to (A) *S. cerevisiae* flocculins Flo1,5,9 and 10 (five ORFs), (B) *S. cerevisiae* flocculin Flo11 (three ORFs), or (C) containing a β-sandwich similar to the structure present in *C. albicans* adhesin Als3 (three ORFs).

Known ASL wall proteins in baker’s yeast and *Candida* spp. are Pir proteins, Bgl2 and Sun4 family proteins (described above), and Tos1. Homologs of these proteins were also identified in *C. krusei* (S1 Table). The Pir protein family has a proposed role in cell wall reinforcement through β-1,3-glucan crosslinking. The *PIR* family in *C. krusei* and *S. cerevisiae* is expanded compared to *C. albicans* (Ck and Sc 5 vs Ca 2 genes).

### The cell wall composition of *C. krusei*

To establish relationships between the biosynthetic machinery as deduced from genomic studies and the abundance of cell wall macromolecules in *C. krusei*, we determined its cell wall composition. First, the amount of glucan and mannan was determined by HPLC analysis after acid hydrolysis of purified walls from exponentially growing cells and biofilms (Fig 2A). In both conditions *C. krusei* contained about 50% glucan (determined as glucose), which is slightly less than the 61.5% in *C. albicans*. In contrast, a significantly higher amount of mannan is present in walls of *C. krusei* (Ck: 40% vs. Ca: 16%), consistent with differences in gene copy numbers of mannosyltransferase protein families discovered in the genomic analysis. In a complementary approach, α-mannan was determined by binding to ConA and phosphomannan by Alcian blue binding. Coherent with the HPLC and genomic data, the flow cytometry (FC) analysis showed elevated levels of ConA binding in *C. krusei* compared to *C. albicans* (Fig 2B), also leading to increased cell aggregation in *C. krusei* but not in *C. albicans* (Fig 2D). In contrast, reduced Alcian blue binding was observed (Fig 2B), consistent with the reduced gene copy number of β-mannosyltransferases in the genome of *C. krusei*.

**Figure 2.**
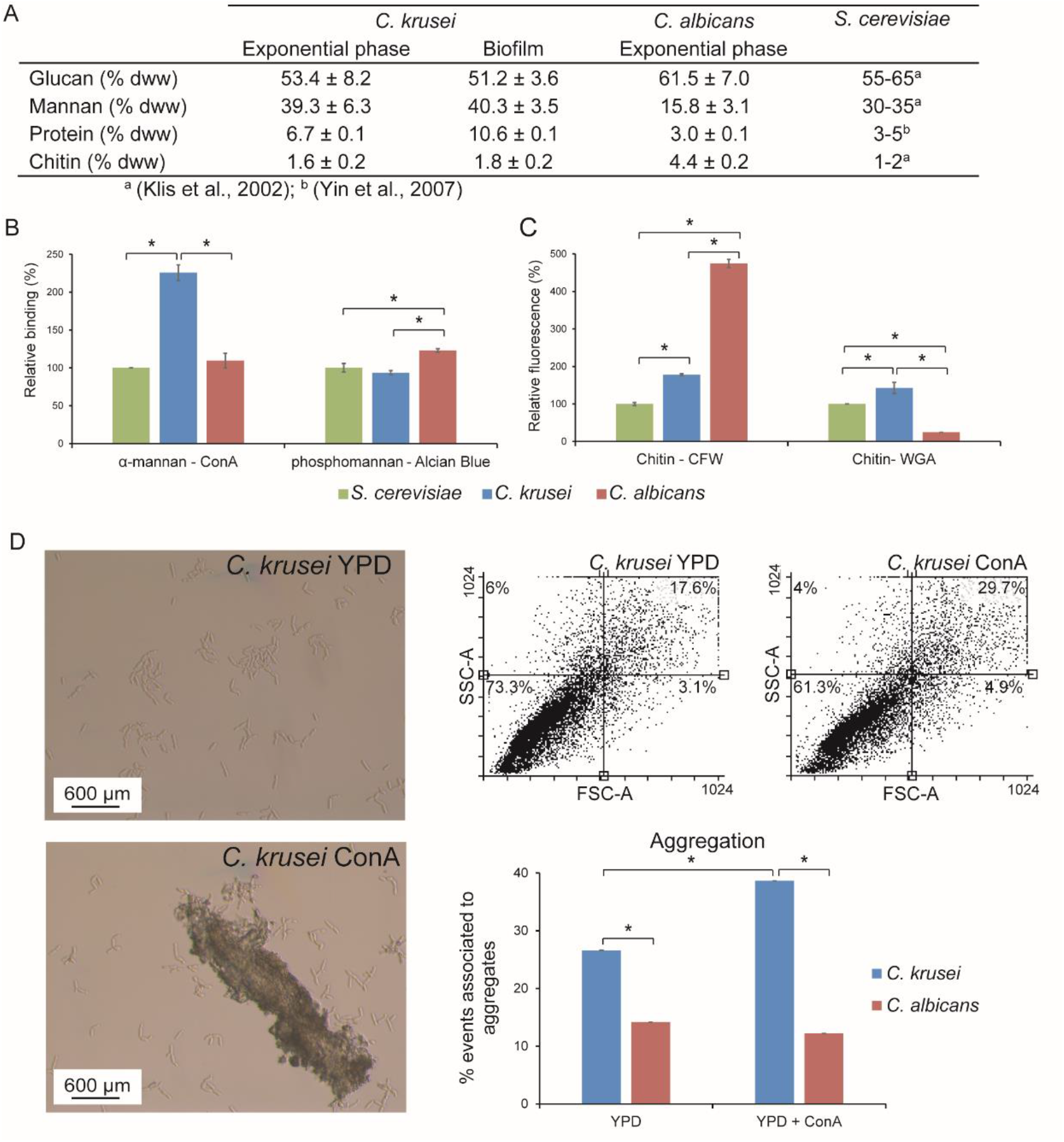
*C. krusei* cell wall composition analysis. (A) Glucan, mannan, protein and chitin levels in the cell wall of *C. krusei* determined upon chemical hydrolysis. (B and C) Flow cytometry (FC) and colorimetric analysis of (B) Concanavalin A (ConA) α-mannan binding and Alcian blue β-mannan binding (C) chitin-binding Calcofluor white (CFW) and Wheat germ agglutinin (WGA). (D) Aggregation. Events associated to aggregates were quantified by flow cytometry, by measuring 10,000 cell particles per strain in the presence/absence of Concanavalin A (ConA), FSC-A, particle size; SSC-A, particle complexity. Percentage of cell particles in each quadrant are indicated, cell aggregation data was supported by optical microscopy. *C. albicans* (A, B, C, and D) and *S. cerevisiae* (B and C) are added for comparative reasons. Data in (B) and (C) are normalized against *S. cerevisiae*.

Protein and chitin levels were determined using colorimetric assays upon alkali and acid hydrolysis, respectively, and further explored using FC of CFW and WGA-FITC binding. Walls from exponentially growing *C. krusei* cells contained 6.7% protein, about twofold higher than the amount in *C. albicans* (Fig 2A).

This further increased to 10.6% in biofilm cell walls (Fig 2A). The determined amount of chitin present in the cell wall of *C. krusei* was lower than in *C. albicans* (Ck/Ca = 0.37) but similar to *S. cerevisiae*, and this was supported by CFW binding (Fig 2A and 2C). On the other hand, while WGA-FITC binding was also similar as in *S. cerevisiae*, it was about fivefold lower in *C. albicans* (Fig 2C), which is possibly related to lowered binding to *N*-acetylglucosamine monomers present in CWPs.

The above-described differences in cell wall composition between *C. krusei* and *C. albicans* may affect cell surface properties. Indeed, compared to *C. albicans, C. krusei* shows reduced sensitivity to zymolyase and Congo red (CR) while its sensitivity to CFW is increased (Fig 3A-B).

**Figure 3.**
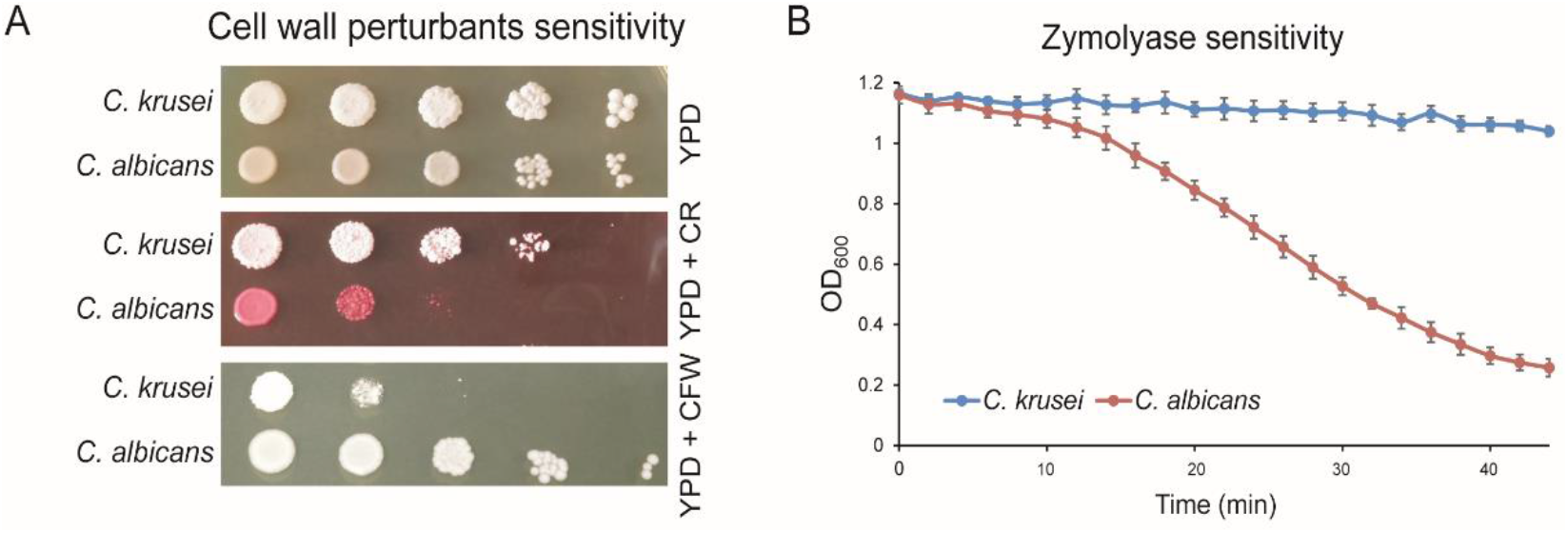
Cell surface properties of *C krusei* compared to *C. albicans*. (A) Spot assays to determine sensitivity to cell wall-perturbing agents Congo red (CR) and Calcofluor white (CFW). (B) Zymolyase sensitivity.

### Floating biofilm formation and adhesion

We then performed experiments to evaluate *C. krusei* biofilm formation and adherence. First, biofilm formation onto polystyrene during 24 h of incubation in YPD without shaking was analyzed. In *C. albicans* this leads to cell sedimentation and subsequent biofilm formation onto the plastic. Interestingly, under these conditions *C. krusei* did not sediment but in contrast formed a film of floating cells, adhered to each other, at the air-liquid interface (Fig 4A). Nevertheless, adhesion of *C. krusei* to polystyrene at the air-liquid interface equaled the biofilm mass of *C. albicans* formed as a layer at the bottom in static cultures (Fig 4C). Aeration by agitation (200 rpm) leads to the formation of *C. krusei* flocs that, compared to *C. albicans*, showed higher adhesion to polystyrene, probably because agitation diminishes biofilm formation of the latter (Fig 4C). Consistent with the observed flocculation (Fig 4B), growth (OD_600_) measurements of liquid cultures showed that *C. krusei* reached a lower maximum cell density (Fig 4B).

**Figure 4.**
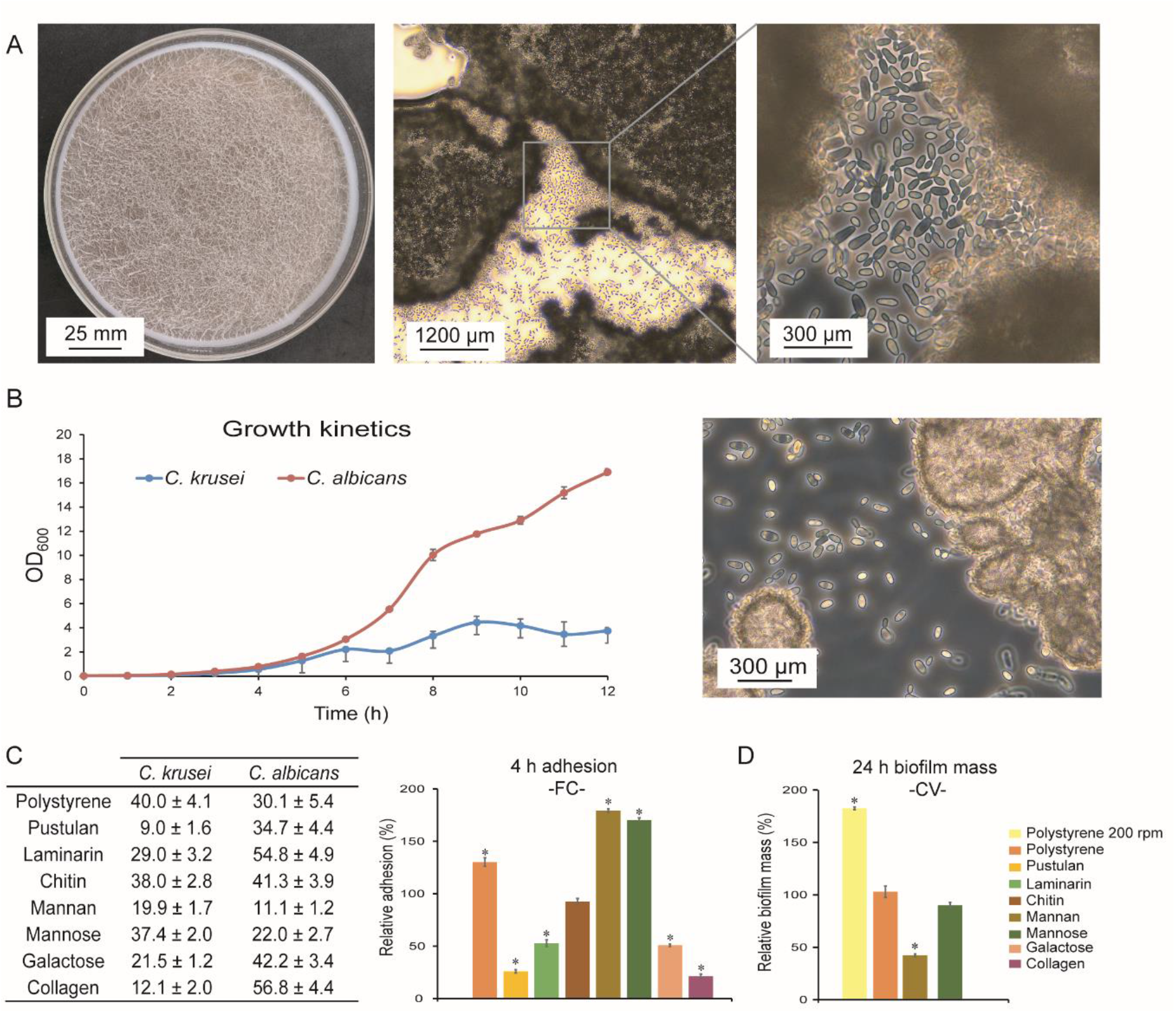
Adhesion and biofilm-forming properties of *C. krusei*. (A) *C. krusei* floating biofilm formation after 24 h static incubation at 37 °C. Left, Petri dish showing floating biofilm; Middle and right, light microscopy images (B) Growth kinetics. On the right, floc formation of *C. krusei* after 12 h. (C) Adhesion to polystyrene and different cell wall and host surface molecules after 4 h of incubation measured by flow cytometry (FC). Biofilm biomass after 24 h measured by Crystal Violet (CV) staining. *C. krusei* data presented in the histograms of (C) and (D) are normalized to *C. albicans*.

Adhesion tests in PBS after 4 h of incubation (without shaking) showed that adhesion to polystyrene was also higher in *C. krusei* than in *C. albicans* (Fig 4C). We then tested whether binding to cell wall components may play a role in the observed cell-cell interactions (Fig 4C). However, binding experiments to immobilized molecules that form the internal polysaccharide layer of the cell wall, pustulan (β-1,6-glucan), laminarin (β-1,3-glucan), and chitin, showed less adhesion of *C. krusei* to these molecules compared to *C. albicans* (Fig 4C). In contrast, a significant increase was observed in binding of *C. krusei* to mannan and mannose (Fig 4C). With regard to host surface molecules, adhesion of *C. krusei* to collagen and galactose was lower than in *C. albicans* (Fig 4C). The presence of mannose did not affect polystyrene-adhered biofilm formed by *C. krusei* but addition of mannan led to decreased biofilm (Fig 4C), again implicating a role for cell wall mannoproteins.

### Proteomic analysis of *C. krusei* flor reveals increased incorporation of adhesins and hydrolytic enzymes

Cell aggregation in *S. cerevisiae* has been shown to be related to the expression of cell wall proteins that act as flocculins (45). Our genomic studies revealed the presence of a similar family of flocculins in *C. krusei*. This prompted us to perform a detailed investigation of the cell wall proteome in *C. krusei*.

Proteomic analysis of cell walls from exponentially growing cultures and 24-h floating biofilms identified a total of 33 genuine CWPs whose precursors either contain signal peptides for secretion or have homology to known CWPs in other species (Fig 5A, S3 Table). All proteins except one were identified in both conditions, the remaining one, Flo10, was identified only in floating biofilms (Fig 5A and S3 Table). Consistent with our previous analyses in other *Candida* spp, the identified proteins can roughly be divided into two groups: (i) core wall proteins with similar abundance in both conditions, and (ii) condition-dependent proteins, mostly adhesins, upregulated during biofilm formation.

**Figure 5.**
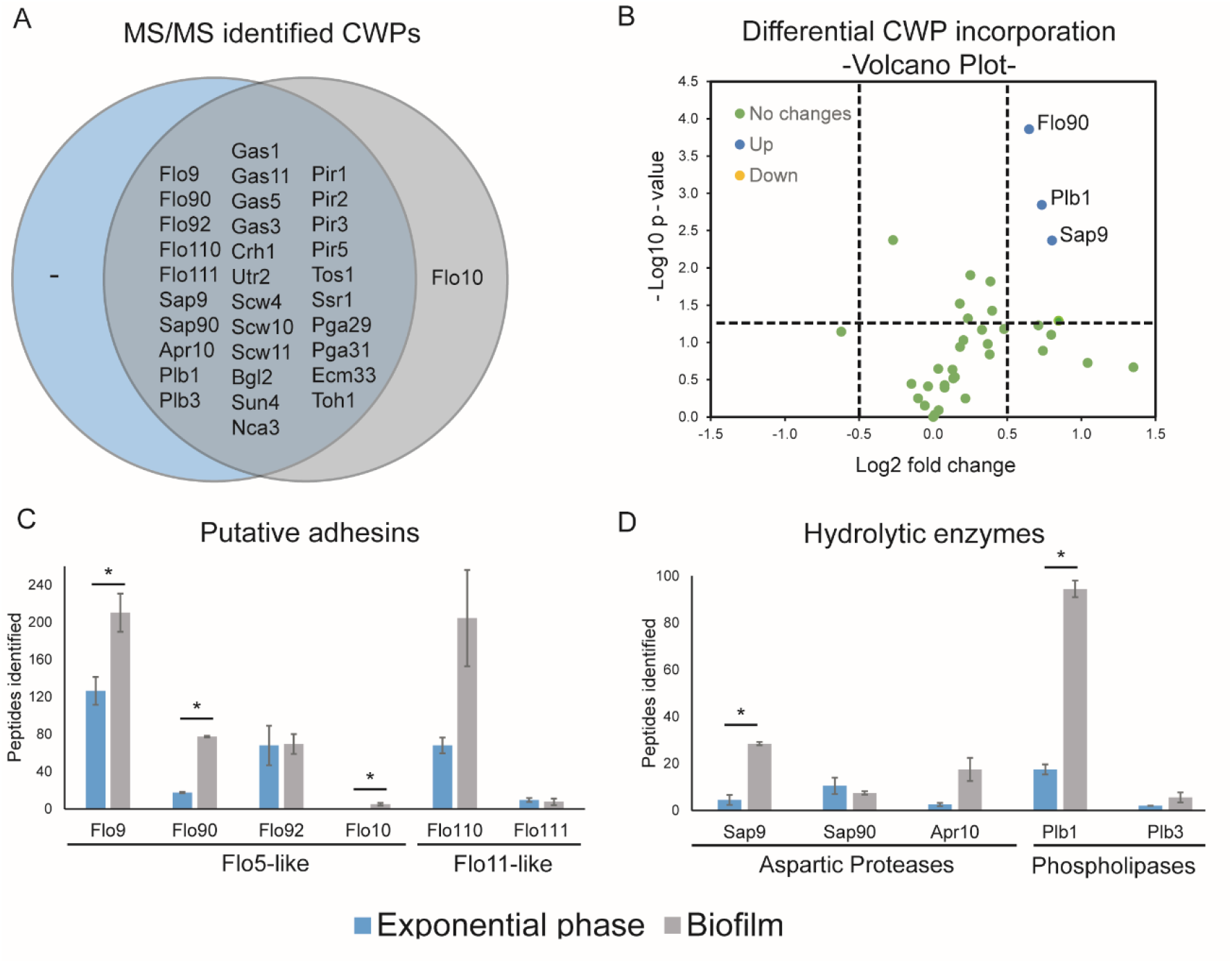
Comparative proteomic analysis of *C. krusei* cell walls. (A) Venn diagram showing comparative analysis of exponential phase cells and biofilms. (B) Analysis of individual protein abundance in biofilms versus exponential phase cells based on normalized peptide counts. Indicated are proteins with at least twofold and statistically significant changes in abundance. Peptide counts of (C) putative adhesins and (D) hydrolytic enzymes. Statistically significant changes (p<0.05) between the two conditions are marked by asterisks.

The identified core wall proteome includes carbohydrate-active enzymes from Crh, Gas, and Bgl2/Scw4 families involved in cell wall polysaccharide remodeling, non-enzymatic Pir and Srp1/Tip1 proteins with proposed roles in crosslinking of β-glucan chains, and proteins with unknown functions including Ssr1 and the widely distributed Ecm33 (Fig 5A).

Semi-quantitative Volcano plot analysis of the proteomic data by peptide counting after normalizing the spectral counts of the core proteome in both conditions (S4 Table), showed elevated incorporation of the adhesin Flo90 as well as the aspartyl protease Sap9, and phospholipase Plb1 in flor biofilms (Fig 5B). Although only Flo90 reached a statistically significant twofold change, four of the six identified putative flocculins appear to have increased abundance in biofilms. Flo9 is abundantly present in flor cells (210 peptides identified) but did not reach a twofold increase, Flo10 was identified only in biofilm (5 peptides). Flo110, was also very abundant in biofilms (205 peptides) and reached a threefold change but lacked statistical significance due to variation between the biological duplicates (Fig 5C and S3 Table). Inferred from its similarity to *S. cerevisiae* Flo11, this protein probably plays an important role in the observed flor formation of *C. krusei*.

With respect to the hydrolytic enzymes found in the cell wall of *C. krusei*, the abundance of aspartyl proteases tripled (18 vs. 54 peptides), while phospholipases almost sextupled (20 vs. 112 peptides) (Fig 5D and S3 Table). Besides the above-mentioned hydrolytic enzymes, abundance in biofilms also appeared increased for Apr10 (protease) and Plb3 (phospholipase), however, in both cases, without reaching statistical significance. This is the first time that such a significant increase of these hydrolytic enzymes has been observed in *Candida* cell walls during biofilm formation, which perhaps might be related to the special characteristics of the floating biofilm produced by this yeast.

## Discussion

The yeast *Candida krusei* (teleomorph *Pichia kudriavzevii*) is among the five most frequent etiological agents of candidiasis but is phylogenetically very distinct to *C. albicans* and other clinically relevant *Candida* species. Perhaps for this reason, *C. krusei* is relatively understudied (4). This is also the case for its cell wall composition even though it is well known that the yeast cell wall plays an important role in the primary host-pathogen interactions that underlie the establishment of infections.

Here, we shed light on the cell wall composition of *C. krusei* through an integrative study including bioinformatic, biochemical, microscopic, and proteomic studies. First, through detailed *in silico* analysis the genetic machinery available to synthesize the cell wall of *C. krusei* is presented. Although our analysis included search queries for enzymes synthesizing cell wall components that have not been described in ascomycetous budding yeasts, such as α-glucan and cellulose (46), only protein families also described for *C. albicans* and/or *S. cerevisiae* yielded positive results, indicating that the composition and general architecture of the cell wall in *C. krusei* is similar to these yeasts. In this respect, our data corroborate the data presented by Navarro-Arias and colleagues (47) who documented the presence of β-1,3-glucan, chitin, *N*- and *O*-linked mannan and phosphomannan in *C. krusei*. Our bioinformatic analysis revealed that the biosynthetic glucan and chitin gene repertoire is very similar as in *C. albicans*, from which one could hypothesize that roughly similar amounts of these polysaccharides could be expected in the cell wall of *C. krusei*. The biochemical analyses mostly support this notion, however, two independent approaches, canonical acid hydrolysis with hot 6 N HCl followed by a colorimetric assay and CFW staining indicate that the level of cell wall chitin is about twofold lower in *C. krusei* than in *C. albicans*. The latter contrasts with Navarro-Arias et al. who documented elevated levels of chitin in *C. krusei* (47). However, their data are relative (and not absolute) values produced by combined HPLC measurements of glucan, mannan, and chitin monomers in cell wall hydrolysates and assumed to account for 100% of the cell wall weight, obviously not a correct assumption. Interestingly, in contrast to CFW, our WGA-FITC binding assays showed increased staining of *C. krusei* compared to *C. albicans*, which was also observed by Navarro-Arias et al. The difference between the CFW and WGA results may be explained by differential binding specificities of these molecules for different types of chitin molecules (48), a more difficult access of the larger molecule WGA to chitin buried in internal layers of lateral walls (49), or to the fact that CFW not only binds chitin but also has some affinity for linear β-1,3-glucan (50).

In contrast to synthesis of glucan and chitin, differences in gene numbers for families related to degradation or modification of these polymers were observed between the genomes of *C. albicans* and *C. krusei*. Lowered numbers of glucanases and chitinases whose orthologs in other species are involved in cytokinesis (51) does not directly imply that cell division is occurring less in *C. krusei* as cell clumping is infrequent under unlimiting growth conditions. However, it is noteworthy that, even under standard laboratory conditions, *C. krusei* cultures tend to show a high percentage of cells that are more elongated (Figs 2 and 4) and can easily be distinguished from cultures with typical oval-shaped *C. albicans* cells.

Strikingly, large differences in some of the gene families responsible for protein mannosylation during secretion were observed between *C. krusei* and *C. albicans*, suggesting major differences in glycosylation structures, especially of *N*-glycans between the two species. More specifically, the extended family of GT32 α-1,6-mannosyltransferases in *C. krusei* suggests it may have longer *N*-glycan α-1,6-mannan outer chain backbones. These backbones may be (more) heavily decorated with short α-1,2-linked branches because of the expansion of GT15 and GT71 α-1,2-mannosyltransferases, which might also result in longer *O*-glycans. On the other hand, reduced Mnn4 (GT34) and Bmt (GT91) gene families suggests a lower level of phosphomannan with fewer additions of β-1,2-linked mannopyranosides to the branches (Fig 6). Our biochemical analyses are consistent with these genomic data; while the mannan content in *C. krusei* is 2.5-fold higher than in *C. albicans*, a reduction in phosphomannan was detected. Of course, we need to keep in mind that the mannan level will be related to the cell wall protein content, which is also twofold higher in *C. krusei* than in *C. albicans*. Interestingly though, where the protein content was further increased in biofilm cells, this was not or hardly the case for the mannan level.

**Figure 6.**
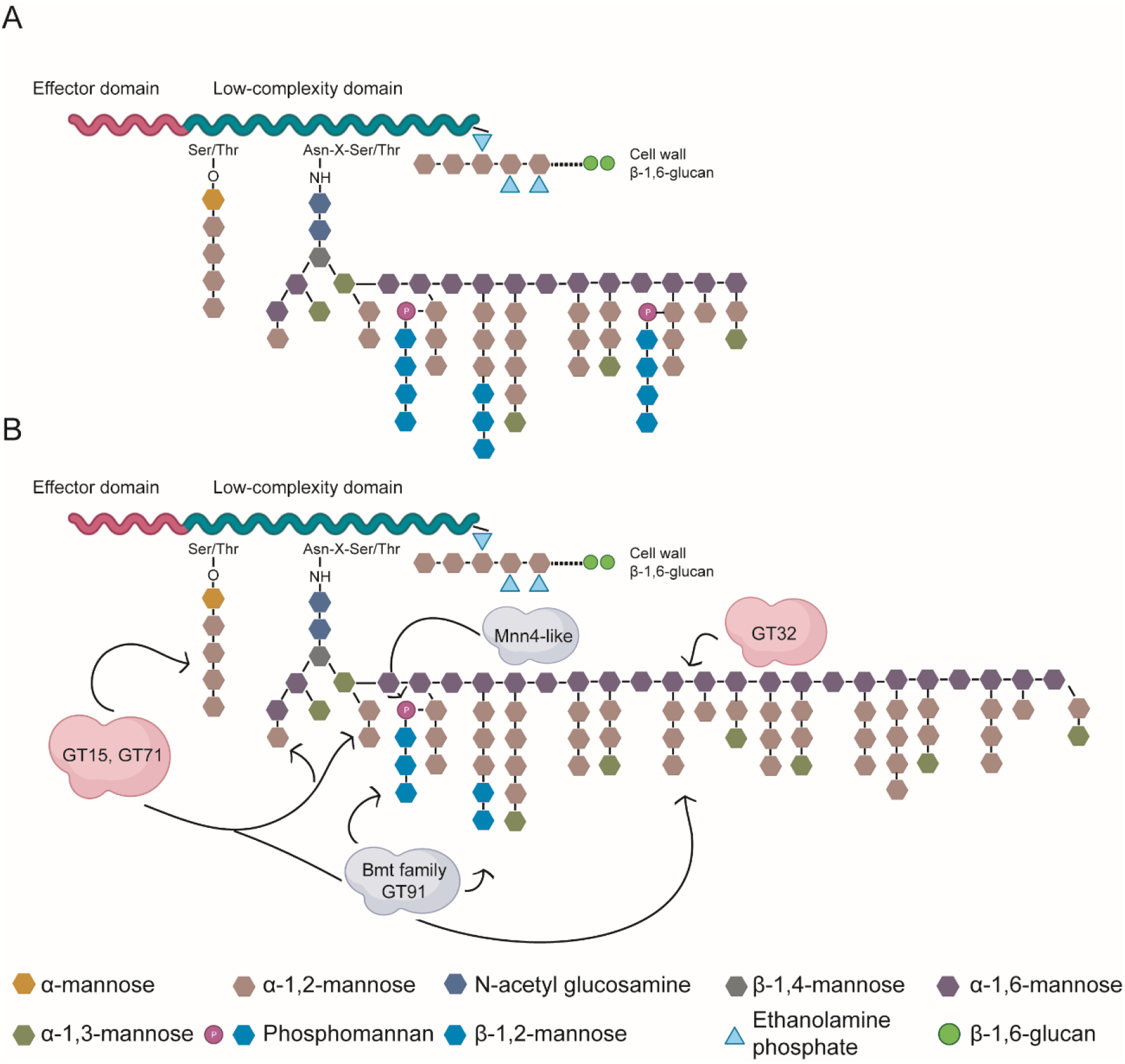
Schematic model showing cell wall protein mannosylation. (A) *C. albicans* protein mannosylation as described (52). (B) Protein mannosylation in *C. krusei* as proposed in this study. Indicated in clouds are protein families expanded (pink) and reduced (grey) compared to *C. albicans*.

Evaluation of biofilm formation and adherence in *C. krusei* led to the formation of floating biofilms and flocs, phenotypes that in *S. cerevisiae* are associated with the expression of flocculins in the cell wall (27). The flocculin (Flo) family of *S. cerevisiae* includes homologous Flo1, Flo5, Flo9 and Flo10 lectins that promote cell-cell adhesion by binding to mannose, which may lead to floc formation. Interestingly, their N-terminal domains, so-called PA14 domains, show structural similarity to those in Epa proteins of the pathogen *C. glabrata*, which have been shown to bind oligosaccharides with terminal galactosyl residues of human glycoproteins, suggesting a direct role in host-pathogen interaction (53, 54). On the other hand, *S. cerevisiae* also contains the apparently unrelated Flo11 that promotes cell-cell adhesion in a weaker manner, contains sequences promoting the formation of amyloids (55) (S1 Fig), and is essentially required for biofilm formation (56). The *C. krusei* genome encodes at least eight putative flocculins, including five Flo5-like and three Flo11-like proteins. Mass spectrometric analysis of purified walls revealed the presence of six of these flocculins (four Flo5-like and two Flo11-like), four of which are upregulated in floating biofilm cells compared to the exponential growth phase, as judged by peptide counting. Presence of peptide sequences with possible roles in β-aggregation was suggested by Tango prediction for both Flo11-like proteins (S1 Fig). Interestingly, Flo110 is one of the most abundant wall proteins in *C. krusei* floating biofilm cells. Altogether, this suggests that these proteins, but especially Flo110, may play a key role in mediating formation of the floating biofilm.

Finally, the increased abundance in biofilm cells of hydrolytic enzymes, the aspartic proteases Sap9 and Apr10 and phospholipases Plb1 and Plb3, fits well with increased Sap9 and Plb5 levels in walls of *C. albicans* cells grown with lactate as carbon source compared to glucose-grown cells (57). Increased Sap9 levels in *C. albicans* cell walls have also been observed under increased temperatures and fluconazole stress (58, 59), and expression of *C. albicans SAP9* has been associated with adhesion and damage of epithelial cells (60).

In conclusion, our integrated study provides important novel insights into the biosynthesis, composition, and architecture of the cell wall of *C. krusei*, including the identification of cell wall proteins that might play key roles in biofilm formation and host-pathogen interactions of this pathogenic yeast.

## Materials and Methods

### *In silico* genomic analysis

*C. krusei* bioinformatic analysis was carried out with the genome sequence of reference strain CBS573, assembly [ASM305444v1](https://www.ncbi.nlm.nih.gov/genome/14557?genome_assembly_id=372140) (5), retrieved from NCBI (https://www.ncbi.nlm.nih.gov). Annotations, functional assignment, and homology studies were performed using a local BLAST tool downloaded from EMBOSS http://emboss.sourceforge.net/, for which known fungal cell-wall related protein sequences from related species, mostly *S. cerevisiae* and *C. albicans*, were used as queries. Predicted homologs were judged by identity levels over the whole sequence (generally >30%) and the presence of known functional domains. Putative GPI proteins were identified using a pipeline approach as described (61). In brief, prediction of C-terminal GPI-modification sites was performed using two complementary approaches: (i) a selective fungal-specific algorithm named Big-PI, available at http://mendel.imp.ac.at/gpi/fungi_server.html (62), and (ii) a more inclusive pattern and composition scanning using the web tool ProFASTA, available at http://www.bioinformatics.nl/tools/profasta/, as detailed in (61). Positive proteins from both approaches were filtered using ProFASTA, and further analyzed to select proteins that contain N-terminal signal peptides but lack internal transmembrane domains using SignalP and TMHMM servers at DTU (https://services.healthtech.dtu.dk/), followed by ProFASTA parsing. As a last step in the pipeline, possible false positives were removed following NCBI-BLAST analysis.

### Protein structural modeling and analysis

Three-dimensional structures of CWPs were modeled using the AlphaFold2 algorithm available at the ColabFold server (63) and subsequently visualized using iCn3D (64). Structure similarity analysis against the proteins in the PDB database was performed using two free available algorithms: the DALI server (65) and PDBeFold, both with a default cut-off for lowest acceptable similarity match of 70% (66). Structural pairwise alignment was executed with a web tool from the RCSB Protein Data Bank available at https://www.rcsb.org/alignment. TANGO was used for prediction of β-aggregation (http://tango.crg.es/about.jsp)

### Cell culturing

Yeast strains used in this study are the reference strains *C. krusei* CBS573, *C. albicans* SC5314 and *S. cerevisiae* CEN.PK113-7D, the latter two for comparative purposes. Culturing was performed in liquid YPD (1% yeast extract, 2% peptone, 2% glucose) at 37 °C and 200 rpm unless mentioned otherwise. For development of *C. krusei* biofilms, cells were precultured overnight in YPD at 37 °C, and then adjusted to a cell density of OD_600_ = 0.5 in fresh YPD. Twenty mL of the cell suspension was pipetted into a polystyrene Petri dish and incubated at 37 °C for 24 h in a humid environment without shaking. Floating biofilm (flor) was recovered from the top of the culture with a spatula, and biofilm cells were resuspended in PBS and collected by centrifuging.

### Cell wall isolation

Cell wall isolation was performed as described by De Groot et al. (2004). Briefly, cells were harvested by centrifugation and washed with cold 10 mM Tris-HCl, pH 7.5. After resuspension in 10 mM Tris-HCl, pH 7.5, the cells were disintegrated with glass beads using a FastPrep-24 instrument (MP Biomedicals) in the presence of a yeast/fungal protease inhibitor cocktail (Sigma). To remove intracellular contaminants and non-covalently associated proteins, the cell walls were washed extensively with 1 M NaCl, incubated twice for 5 min at 100 °C in a buffer containing 2% SDS, 150 mM NaCl, 100 mM Na-EDTA, 100 mM β-mercaptoethanol and 50 mM Tris-HCl, pH 7.8, and afterwards washed five times with water, each step followed by centrifugation (5 min at 3300 g). Purified walls were lyophilized and stored at -80 °C until use.

### Cell wall components content determination

For the determination of total cell wall glucan and mannan, purified cell walls were hydrolyzed with sulfuric acid (67) and the amount of glucose and mannose was determined by HPLC according as described (68). Values shown are the mean of duplicate measurements from two independent experiments.

Protein and chitin content were determined using colorimetric assays as described by Kapteyn et al. (69). Similarly, the phosphomannan content was also determined colorimetrically using the Alcian Blue (Sigma) binding assay as described by Li et al. (2009) (70).

Mannoproteins were fluorescently stained with Rhodamine-ConA (Fisher Scientific, 100 µg/ mL). Chitin was stained with CFW (Glentham Life Science, 25 µg/ mL) and Wheat germ agglutinin (WGA)-FITC (Sigma, 100 µg/ mL). Fluorescent staining was performed with exponential phase cultures. The fluorescence intensity of the different samples was determined by flow cytometry.

### Aggregation

Cell aggregation of log phase cultures in YPD was analyzed with a Leica DM1000 microscope mounted with a MC170 HD digital camera and by flow cytometry by measuring size (FSC-A) and granularity (SCC-A) of 10,000 cell particles. Effects on the addition of Concanavalin A (ConA) on aggregate formation were studied by incubating log phase cultures for at least 45 min at 37 °C.

### Congo red and Calcofluor white sensitivity spot assays

To reveal possible alterations in cell wall organization, sensitivity to the cell wall-perturbing compounds Congo red (CR) and Calcofluor white (CFW) was assayed using drop tests. Overnight precultures were diluted to an OD_600_ = 1. From these, tenfold dilution series were prepared, and four μL aliquots were spotted on YPD plates containing different concentrations of CFW (50 and 100 μg/mL) and CR (100 and 200 μg/mL). Growth was monitored after 24 h of incubation at 37 °C.

### Zymolyase sensitivity assay

Cells were grown to exponential phase (OD_600_ = 1), centrifuged, and resuspended in 10 mM Tris-HCl, pH 7.5 at an OD_600_ = 2. Then, 2.5 μL/mL β-mercaptoethanol was added, and the cells were incubated for 1 h at room temperature. After this, 1.8 mL of the cell suspension and 200 μL of up to 100 U/mL zymolyase 20T were mixed, and decrease in OD_600_ because of cell lysis was measured every two min after a short mixing over a period of 45 min.

### Quantitation of biofilm formation

For analysis of the biofilm formation capacity, the OD_600_ of overnight precultures was adjusted to 2, and polystyrene 96-wells plates were filled with 50 μL of the cell suspension and 150 μL of fresh YPD medium. After a 24 h incubation in a humid environment with or without shaking, the spent medium was removed, and the biofilms were washed once by immersion in a mQ water bath to remove unbound cells. Biofilm cells were stained with a 0.1% crystal violet (CV) (Sigma-Aldrich) solution for 30 min. Excess of CV was removed, and the cells were washed two times with mQ water. Stained adhered cells were resuspended in 33% acetic acid, and the intensity of CV staining was measured as OD_595_ using a microtiter plate reader SpectraMax 340PC (Molecular Devices, CA).

### Measurement of 4 h adhesion to polystyrene

For the analysis of the capacity to adhere (rather than form biofilms) to polystyrene, overnight precultures of strains of interest were adjusted to 10^6^ cells/mL in PBS, and polystyrene 12-wells plates were filled with 500 μL of the cell suspension. After a 4 h incubation at 37 °C, unbound cells were removed by two washes with PBS. Adhered cells were then disentangled by enzymatic digestion using 100 μL of a 2.5% trypsin (Sigma-Aldrich) solution and resuspended in 500 μL PBS. The quantity of adhered cells was measured using a MacsQuant flow cytometer (Miltenyi Biotec, Madrid, España).

### Adhesion to fungal cell wall components and host surface molecules

Differences in the binding capacity to fungal cell wall components or to the mammalian extracellular matrix (ECM) molecules collagen and galactose was evaluated by flow cytometry. Twelve-wells microtiter plates were coated with pustulan (β-1,6-glucan, Calbiochem; 250 µg/mL in 50 mM potassium acetate buffer), laminarin (β-1,3-glucan, Thermo Fisher; 250 µg/ mL in milli-Q water), chitin (Sigma; 250 µg/mL in 1% acetic acid), mannan (mannan from *S. cerevisiae*, Sigma; 250 µg/ml in milli-Q water), mannose (D- (+)-mannose, Sigma; 250 µg/ml in milli-Q water), galactose (D-(+)-galactose LS, Panreac; 250 µg/ml in milli-Q water) or bovine collagen (Sigma; 250 µg/mL in 0.2 M bicarbonate buffer, pH 9.6) by adding 500 µL of the respective solutions, and allowing for evaporation (overnight at 37 °C), in the case of laminarin, pustulan, mannan, mannose, and galactose, or passive adsorption (1 h at 30 °C followed by overnight incubation at 4 °C), in the case of chitin and collagen. The plates were then washed with PBS. Overnight cultures of strains of interest were adjusted to 10^6^ cells/mL in PBS, and coated wells were filled with 500 μL of the cell suspension. After 4 h of incubation at 37 °C, unbound cells were aspirated and further removed by two washes with PBS. Adhered cells were loosened with a 2.5% trypsin solution, resuspended in 500 μL PBS, and measured using a MacsQuant flow cytometer.

### Growth kinetics

Cells from overnight precultures were diluted to an OD_600_ = 0.1 in fresh YPD and incubated at 37 °C and 200 rpm for 12 h, measuring the OD_600_ every hour. Cell aggregation was determined at the end of the assay by optical microscopy.

### Mass spectrometric identification of CWPs

Sample preparation. Reduction, S-alkylation, and proteolytic digestion of cell walls with Trypsin Gold (Promega, Madrid, Spain) were performed as described (71). Released peptides were freeze-dried and taken up in 50% acetonitrile (ACN) and 2% formic acid. Samples were diluted with a 0.1% trifluoroacetic acid (TFA) solution to reach a peptide concentration of about 250–500 fmol/μL. Duplicate biological samples were analyzed by LC-MS/MS using a timsTOF Pro (trapped ion mobility spectrometry coupled with quadrupole time of flight Pro) mass spectrometer (Bruker Daltonics) equipped with an Ultimate 3000 RSLC nano ultra-high-performance liquid chromatography (UHPLC system) (Thermo Scientific). Two hundred ng of a tryptic digest cleaned on a TT2 TopTips column was injected into a C18 Aurora column (IonOpticks) (25-cm length by 75-μm inner diameter, 1.6-μm particle size). The peptides were eluted from the column by applying a gradient, from 0.1% formic acid 3% ACN to 0.1% formic acid 85% ACN (flow rate, 400 nl/min), in 140 min. For the acquisition cycle, scans were acquired with a total cycle time of 1.16 s. MS and MS/MS spectra were recorded from 100 to 1,700 m/z.

MS/MS database searching. Raw MS/MS data were processed with Data Analysis software (Bruker, Billerica, MA, USA). Resulting .mgf data files were used for searching with licensed Mascot software (Version 2.5.1) against a non-redundant *Candida krusei* protein database containing protein sequences downloaded from NCBI. Simultaneously, searches were performed against a common contaminants database (compiled by the Max Planck Institute of Biochemistry, Martinsried, Germany) to minimize false identifications. Mascot search parameters were: a fixed modification of carbamidomethylated cysteine, variable modification of oxidized methionine, deamidated asparagine or glutamine, trypsin with the allowance of one missed cleavage, peptide charge state +2, +3 and +4, and decoy database activated. Peptide and MS/MS mass error tolerances were 0.3 Da. Probability-based MASCOT scores (http://www.matrixscience.com/, accessed on 15 February 2021) were used to evaluate the protein identifications with a 1% false discovery rate as the output threshold. Peptides with scores lower than 20 were ignored. Unmatched peptides were subjected to a second Mascot search with semitrypsin as the protease setting, N/Q deamidation as extra variable modification, and a cutoff peptide ion score of 40. Semitryptic peptides identified in the second search served solely to extend the sequence coverage of proteins identified in the first trypsin search. Protein identifications based on a single peptide match were only taken into consideration if identified in multiple samples and at least one time in duplicate samples. The total number of peptides (TP) identified for each protein was determined by adding up all MS/MS fragmentation spectra, leading to protein identification in the two duplicate samples.

#### Protein quantitation

Relative abundance of identified CWPs in biofilms and exponentially growing cells was determined by their total spectral MS/MS counts. Proteins were considered to have differential abundance when an at least twofold statistically significant change is detected between biofilms and exponentially growing cells.

### Statistical analysis

Each assay was performed with at least two biological and two technical replicates unless otherwise stated. Statistical analysis was performed using SPSS v21.0 (IBM). Distribution of the data was analyzed by means of a Shapiro-Wilk test. When data showed a normal distribution, parametric Student’s t-test (if two samples), or ANOVA (if > two samples) followed by post hoc Delayed Matching to Sample (DMS) tests were applied. Data showing non-parametric distribution were analyzed with Mann-Whitney’s U-test (if two samples), or Kruskal-Wallis (if > two samples) followed by pairwise U-test comparison. P values <0.05 were considered statistically significant

## Acknowledgments

We thank Henk L. Dekker (Mass Spectrometry of Biomolecules, University of Amsterdam) and Dr. Sergi Ferrer (University of Valencia) for technical support in the analysis of the cell wall protein and sugar composition, respectively.

